# Active-like structure of the ligand-binding domain of GluK2 with L-glutamate and the positive allosteric modulator BPAM344

**DOI:** 10.1101/2023.11.02.565309

**Authors:** Yasmin Bay, Mie Egeberg Jeppesen, Karla Frydenvang, Pierre Francotte, Bernard Pirotte, Darryl S. Pickering, Anders Skov Kristensen, Jette Sandholm Kastrup

## Abstract

Kainate receptors belong to the family of ionotropic glutamate receptors and contribute to the majority of fast excitatory neurotransmission. Consequently, they also play a role in brain diseases. Therefore, understanding how these receptors can be modulated is of importance. Our study provides a dimeric crystal structure of the ligand-binding domain of the kainate receptor GluK2 in complex with L-glutamate and the small molecule positive allosteric modulator, BPAM344, in an active-like conformation. The role of Thr535 and Gln786 for modulation of GluK2 by BPAM344 was investigated using a calcium-sensitive fluorescence-based assay on transiently transfected cells expressing GluK2 and mutants hereof. This study may aid design of tool compounds targeting kainate receptors, elucidating their potential as targets for treatment of brain diseases.

---

In the mammalian brain, the major excitatory neurotransmitter L-glutamate activates G protein-coupled metabotropic glutamate receptors and fast acting ionotropic glutamate receptors (iGluRs). Kainate receptors (KARs) belong to the class of iGluRs. Besides the KAR family, iGluRs comprise three other families: α-amino-3-hydroxy-5-methylisoxazole-4-propionic acid (AMPA) receptors (AMPARs), N-methyl-D-aspartic acid (NMDA) receptors, and Delta receptors. [1]. An imbalance in the glutamatergic system has been shown to be involved in several brain diseases. Specifically, the KARs have been reported to play a role in e.g., epilepsy, depression, and schizophrenia [2], making the understanding of the molecular function of these receptors essential for unraveling their potential as targets for treatment with drugs. Therefore, it is important to investigate how these receptors can be modulated by small molecule ligands.

The KARs can be further subdivided into subtypes assembled from five subunits (GluK1-5). GluK1-3 can form both homomeric and heteromeric receptors, whereas GluK4-5 can form heteromeric receptors only together with GluK1-3 [3, 1]. Intriguingly, amongst GluK1-3, GluK2 is the most abundant KAR subtype in the brain [4]. However, little is known on how the function of GluK2 can be modulated by small molecules.

A common structural topology exists among the four iGluR families [5]. Four subunits from the same receptor family come together in the formation of a tetrameric receptor. A single subunit is composed of a large extracellular part made up of the amino-terminal domain (ATD) positioned on top of the ligand-binding domain (LBD) harboring the binding site for L-glutamate. The LBD has a clamshell-like bi-lobed structure where L-glutamate binds within the cleft between the upper lobe (known as D1) and the lower lobe (known as D2).

The LBD is further connected to the transmembrane domain (TMD) localized within the cell membrane and forming the ion channel with the pore in the middle. The ion channel becomes permeable to cations upon activation. Finally, a carboxy-terminal domain (CTD) resides inside the cell. The formation of a dimer-of-dimers arrangement creates a dimer interface between two LBDs positioned back-to-back that harbors the binding sites for positive allosteric modulators (PAMs).

A study by Larsen et al. [6] reported BPAM344, a small molecule PAM to possess activity on recombinant KARs despite being a powerful AMPAR PAM. Although enhancing KAR currents 15-fold at GluK2 compared to 5-fold at AMPARs (GluA1i), the potency at GluK2 receptors was determined to be in the micromolar range (EC50 79 µM) [6] and nearly 90-fold lower compared to the AMPAR GluA2 (EC50 0.9 µM) [7].

BPAM344 was first co-crystallized with the LBD of a flip-like AMPAR (GluA2-LBD L504Y-N775S) in the presence of L-glutamate and the structure determined by X-ray crystallography [7], showing a distinct binding mode of two BPAM344 molecules at the dimer interface. Later, the crystal structure of GluK1-LBD in complex with kainate and BPAM344 was determined [6] and showed BPAM344 to adopt a binding mode similar as in the GluA2-LBD structure. Recently, cryo-electron microscopy (cryo-EM) structures were determined of the full-length GluK2 in a closed state in complex with BPAM344 [8].

However, a structure of full-length GluK2 in complex with both BPAM344 and L-glutamate could not be obtained [8]. Here, we report the structure of an active-like dimer of the ligand-binding domain of GluK2 (GluK2-LBD) crystallized with L-glutamate and BPAM344. The structure was determined at 1.6 Å resolution. To explore the importance of two GluK2 residues (Thr535 and Gln786) in the BPAM344 binding site that differ between GluK2 and GluA2, we created two single and one double GluK2 mutants with the corresponding residues in GluA2 (T535S and Q786S). The functional implications were investigated using a calcium-sensitive fluorescence-based assay.

## Materials and methods

### GluK2-LBD for structure determination with L-glutamate and BPAM344

A recombinant construct of GluK2-LBD from *Rattus norvegicus* comprising residues 430-544 from segment S1 and residues 667-805 from segment S2 was used. The two segments were combined with a GT linker as previously reported [9]. A purification tag consisting of six histidine residues and trypsin cleavage site was inserted at the N-terminus. GluK2-LBD was successfully expressed in *Escherichia coli* (*E. coli*) Origami^TM^ 2 cells and the protein purified by His-tag affinity chromatography, ion-exchange chromatography, and size exclusion chromatography. The purity was confirmed by SDS-PAGE (data not shown) and the purified protein used for crystallization employing the hanging-drop vapor diffusion method.

### Protein expression

The pET-28a plasmid expressing the GluK2-LBD construct was transformed into competent Origami^TM^ 2 cells (Novagen). Cells were plated out on a Lysogeny Broth (LB, Invitrogen) agar plate supplemented with the respective selective markers, 30 µg/mL of kanamycin and 12.5 µg/mL of tetracycline (Sigma-Aldrich). A single bacterial colony from the successful transformation was inoculated with 100 mL of LB media overnight at 37°C, afterwards cultured in 1 L Hyper Broth^TM^ (Athena AS) supplemented with glucose nutrient, and the selective markers incubated for 5-6 hours at 37°C. Cells were grown until an OD600 of ∼1.2 was reached and expression induced by IPTG (0.2 mM) and further grown at 25°C overnight with agitation.

### Protein purification

The bacteria culture was harvested by centrifugation and the pellet resuspended in lysis buffer (50 mM sodium phosphate, 10 mM sodium chloride, 1 mM magnesium chloride, 0.06 mg/mL DNAseI, 0.2 mg/mL lysozyme, 0.6 µM PMSF, and 1 EDTA-free protease inhibitor tablet per 50 mL lysis buffer (Roche), pH 7.4). The solution was run through a cell disruptor twice, and collections of the flowthrough were centrifuged prior to loading the solution onto a HisTrap^TM^ HP column (GE Healthcare). The column was washed with buffer 1A (20 mM imidazole, 0.5 M sodium chloride, 50 mM sodium phosphate, pH 7.4) and the protein eluted with a gradient (0-80%) of buffer 1B (0.5 M imidazole, 0.5 M sodium chloride, 50 mM sodium phosphate, pH 7.4). Pooled fractions containing GluK2-LBD were dialyzed against buffer 2A (20 mM sodium acetate, 1 mM L-glutamic acid, 10 mM EDTA, pH 5.0). The tag was cleaved off with trypsin (0.4 mg), and afterwards pH of the solution was adjusted to 5.0 and PMSF added. The solution was loaded on a HiTrap SP HP cation-exchange column (GE Healthcare). The protein was eluted with a gradient (0-30%) of buffer 2B (20 mM sodium acetate, 1 mM L-glutamic acid, 10 mM EDTA, 0.5 M sodium chloride, pH 5.0). Pooled fractions containing GluK2-LBD were then dialyzed against crystallization buffer (10 mM HEPES, 20 mM sodium chloride, 1 mM EDTA, pH 7.0) at 4°C using a dialysis bag. Finally, the protein solution was concentrated and further purified utilizing size exclusion chromatography with a Superdex^TM^ 75 column (GE Healthcare). The protein solution was concentrated to 9.2 mg/mL and used for crystallization.

### Crystallization

BPAM344 was dissolved in DMSO and L-glutamic acid in crystallization buffer. BPAM344 and L-glutamic acid were added to the protein solution, resulting in a final concentration of 10 mM BPAM344, 8.8 mM L-glutamate, and 5.1 mg/mL GluK2-LBD (protein mix). A crystallization screen was designed inspired by Nayeem et al. [10]: PEG4000 ranging from 10-30%, propan-2-ol from 3-12%, 0.1 M sodium acetate pH 5.5, and with or without 0.12-0.24 M sodium chloride. 1 µL of the protein mix was mixed with 1 µL of reservoir solution from the crystallization screen on a siliconized glass cover and inverted over a well of a 24-well VDX plate (Hampton Research). The crystal used for data collection was obtained at room temperature (r.t.) and with reservoir solution consisting of 25% PEG4000, 9% propan-2-ol, 0.1 mM sodium acetate, pH 5.5. The crystals were cryoprotected using reservoir solution supplemented with 20% glycerol prior to flash cooling in liquid nitrogen. X-ray diffraction data was collected at the BioMAX beamline, MAX IV Laboratory in Lund in Sweden [11].

### Structure determination

The diffraction data was processed in XDS [12] and Scala [13] within the CCP4i software [14]. Molecular replacement was performed in Phaser [15] within CCP4i, using the GluK2-LBD structure in complex with L-glutamate (PDB ID: 2XXR) [10] as a search model. The obtained solution was used to build a model with Autobuild in Phenix [16]. Ligands, ions, and missing residues were manually built-in using Coot [17] and inspection of the structure was performed in-between refinement rounds in Phenix. For refinements in Phenix, Translation-Libration-Screw (TLS) groups were defined, and individual B values and riding hydrogen atoms applied. Generation of coordinate file for BPAM344 was performed using the program Maestro (Release 2021-3, Shrödinger, LLC, New York, NY, 2021). The BPAM344 parameter file was acquired from eLBOW [18], maintaining the geometry obtained from geometry optimization in MacroModel (Maestro Release 2021-3, Schrödinger, LLC, New York, NY, 2021) and used for refinements in Phenix. The DynDom server [19] was used to estimate domain closures of GluK2-LBDs relative to the apo structure of the LBD from the full-length GluK2 structure (PDB ID: 8FWQ, chain A) [8]. At last, the structure was validated in the wwPDB OneDep [20] and figures were prepared in PyMOL (The PyMOL Molecular Graphics System, Version 2.5.4 Schrödinger, LLC).

### Molecular Biology

Plasmid DNA containing rat WT GluK2 cDNA in the unedited (*Q*) form (pcDNA3-GluK2(*Q*)2a) was generously provided from Dr. C. Mulle (University of Bordeaux, France). Transfection-grade DNA was purified from *E*. coli bacterial cultures using NucleoSpin Plasmid QuickPure or NucleoBond Xtra MidiPrep kits (Macherey-Nagel, Düren, Germany) according to manufactureŕs instructions. Generation of GluK2 point mutants was performed by site-directed mutagenesis using the QuickChange Mutagenesis Kit (Stratagene, La Jolla, CA). Forward and reverse primer for creation of mutants were: T535S_for: CAAAGCCGTTTATGTCACTTGGAATAAG; T535S_rev: TTATTCCAAGTGACATAAACGGCTTTG; Q786S_for: CTTCAGCTGTCGGAGGAAGGC; Q786S_rev GCCTTCCTCCGACAGCTGAAG. All mutants and WT cDNA constructs for GluK2 were verified by sequencing of the full coding region (GATC Biotech). The WT GluA2 construct carrying the unedited (*Q*) flip isoform (rat GluA2(*Q*)i-pXOOF) was kindly provided by Dr. C. Stenum-Berg (University of Copenhagen) and generated as described in Stenum-Berg et al. [21].

### Culturing of cells

GT-HEK293 cells were cultured in monolayer in a humidified 5% CO2 atmosphere at 37°C in Dulbeccós Modified Eagle Medium (DMEM) with GlutaMAX, supplemented with 10% fetal bovine serum, 100 units/mL penicillin, 100 µg/mL streptomycin, and 1% geneticin (G418) (Sigma-Aldrich). Un-transfected cells were passed by trypsination every second to third day and generally maintained until passage thirty. For plate-based assays, the GT-HEK293 cells were transiently transfected in suspension using LipoD293 (SignaGen) transfection reagent according to manufactureŕs guideline. For each well in a 96-well plate, the transfection mix contained 0.1 µg DNA in a 1:3:10 ratio with LipoD293 and DMEM.

### Calcium-sensitive fluorescence-based assay

Cells for the detection of intracellular calcium were grown on Poly-*D*-Lysine coated black clear-bottom 96-well plates (GR-655087, Greiner, Germany) for at least 48 hours after transfection and reached a cell confluence of around 70-90%. On the day of conducting the experiments, transfected cells were washed two times in phosphate bovine serum (PBS) buffer and loaded with a FluoAM-8 dye (AAT Bioquest) solution in plain DMEM (r.t.), reaching a final concentration of dye per well of at least 2 µM. Loaded cells were incubated at 37°C for at least 30 minutes and excess dye was washed off by PBS, re-applied three times prior to incubating the cells with 50 µL per well assay buffer (100 mM choline chloride, 10 mM sodium chloride, 5 mM potassium chloride, 1 mM magnesium chloride, 20 mM calcium chloride, 10 mM HEPES, pH 7.4). Drug compound solutions were prepared in assay buffer in clear V-bottom 96-well plates (GR-651161, Greiner). Typically, experiments were performed in triplicate or quadruplicate wells for each construct and repeated on 3-4 independent days. Recordings of FluoAM-8 fluorescence were performed using FlexStation I plate reader (Molecular Devices) with an excitation wavelength of 485 nm and detection of emission at a wavelength of 538 nm over a total period of 120 seconds per well. Following a baseline measurement of 16 seconds, receptor induced changes in intracellular calcium concentration were continuously detected after automated addition of the compound solution by the FlexStation machine using black pipette tips (Molecular Devices).

### Data analysis

FlexStation data was analyzed using SoftMax Pro version 5.4 (Molecular Devices) to quantify peak fluorescence response to agonist application in presence and absence of PAM. Specifically, peak fluorescence was calculated as the difference between maximal observed agonist-evoked increase in fluorescence and pre-agonist baseline fluorescence. For construction of concentration-response curves, peak fluorescence from tri- or quadruplicate wells representing identical PAM and agonist concentrations were averaged and normalized to the maximum average responses in the individual plate experiment. Composite concentration-response curves were generated by plotting normalized data from multiple experiments as a function of PAM or agonist concentration using GraphPad Prism software and fitted to a four-parameter Hill equation:

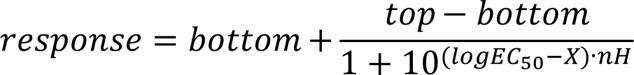

where *bottom* is the fitted minimum response, *top* is the fitted maximum response, *nH* is the Hill slope, and *X* is the concentration, and *EC50* is the half-maximally effective concentration of PAM.

## Results and Discussion

### X-ray structure of GluK2-LBD with L-glutamate and BPAM344

The structure of GluK2-LBD in complex with L-glutamate and BPAM344 was determined at 1.6 Å resolution (Table 1), unraveling the binding mode of ligands and ions at GluK2 in an active-like conformation. GluK2-LBD crystallized as two dimers, with one dimer made up of chain A and a symmetry-related chain A (chain Asym) (Fig. 1A) and the other dimer of chain B and chain Bsym, resembling the dimer previously reported for the structure of GluK2-LBD with L-glutamate only (PDB ID: 2XXR) [10].

**Fig. 1.**
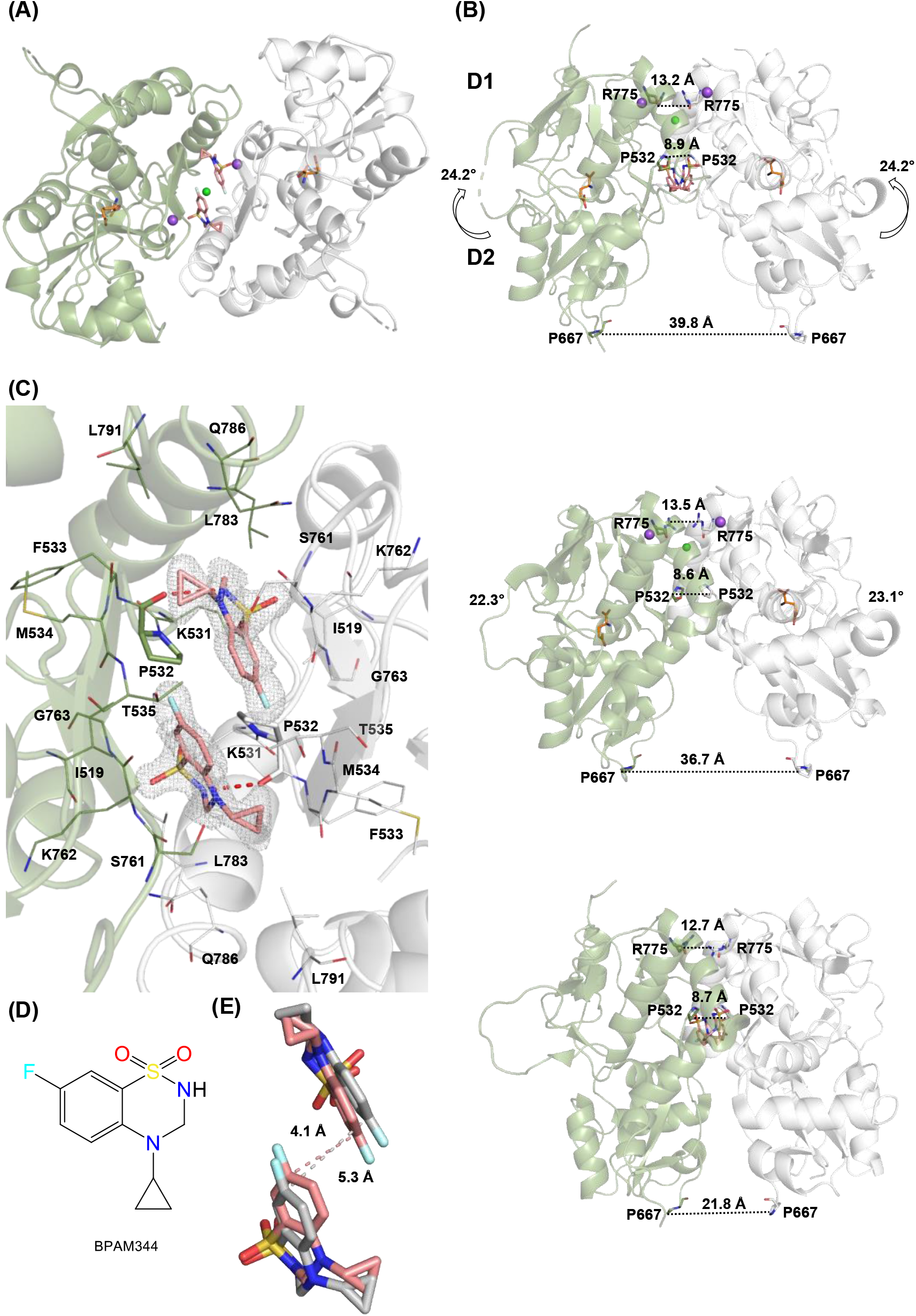
X-ray structure of GluK2-LBD in complex with L-glutamate and BPAM344. (A) Bottom view of the dimeric GluK2-LBD structure (chain A in grey and chain A_sym_ in green). L-Glutamate located at the orthosteric site is shown as sticks with carbon atoms in orange. Two BPAM344 molecules located at the dimer interface are shown as sticks with carbon atoms in salmon. Two sodium ions and one chloride ion also located at the dimer interface are represented as spheres in purple and green, respectively. Oxygen atoms are shown in red, nitrogen atoms in blue, sulfur atoms in yellow, and fluorine atoms in light blue (color scheme used throughout the paper). (B) Upper figure: Sideview of the dimeric GluK2-LBD structure. Distances measured from Pro532, Pro667, and Arg775 in chain A to their respective residues in chain A_sym_ are illustrated as dotted black lines. The arrows indicate the domain closure of lobe D2 towards lobe D1 relative to the LBD (chain A) of the full-length GluK2 apo structure with BPAM344 (lower figure). Middle figure: Sideview of dimeric GluK2-LBD in complex with L-Glutamate (PDB ID: 2XXR) (chain A and B). Lower figure: LBD dimer (chain A and chain D) from the GluK2 full-length cryo-EM structure (PDB ID: 8FWQ) in complex with BPAM344. (C) Close-up on the binding site of BPAM344, showing the 2Fo-Fc omit electron density map contoured at 1σ and carved at 1.6 Å. Residues within 4 Å of the two BPAM344 molecules are shown in lines representation. Pro532 in each subunit is shown in sticks and forms a hydrogen bond to BPAM344 (red dashed lines). (D) Chemical structure of the positive allosteric modulator BPAM344. (E) Close-up on BPAM344 molecules from the GluK2-LBD structure in complex with L-glutamate and BPAM344 overlayed with the BPAM344 molecules from the full-length structure of apo GluK2 in complex with BPAM344. The two structures were aligned on lobe D1 residues (chain A). Protein residues have been omitted for clarity. The distance between the two modulators from the same structure is indicated: GluK2-LBD with L-glutamate and BPAM344 as salmon dashed lines and full-length GluK2 with BPAM344 as grey dashed lines.

**Table 1.** Data collection and refinement statistics for the crystal structure of GluK2-LBD in complex with L-glutamate and BPAM344.

|  |  |
| --- | --- |
| **Crystal data** |  |
| Space group | *P*2<sub>1</sub>2<sub>1</sub>2 |
| Unit cell axes *a*, *b*, *c* (Å) | 98.2, 100.7, 48.2 |
| Unit cell axes $\alpha$, $\beta$, $\gamma$ (°) | 90, 90, 90 |
| Molecules in a.u. | 2<sup>a</sup> |
| **Data collection** |  |
| Beamline | BioMAX MAX IV, Lund, Sweden |
| Wavelength (Å) | 0.976 |
| Resolution (Å) | 48.16-1.60 (1.63-1.60)<sup>b</sup> |
| No. of unique reflections | 63898 (3075)<sup>b</sup> |
| Average multiplicity | 13.6 (13.6)<sup>b</sup> |
| Completeness (%) | 100.0 (99.9)<sup>b</sup> |
| *R*<sub>merge</sub> (%) | 10.9 (76.1)<sup>b</sup> |
| Mean *I*/σ(*I*) | 16.3 (3.9)<sup>b</sup> |
| Wilson B (Å<sup>2</sup>) | 10.6 |
| CC<sub>1/2</sub> | 1.00 (0.91)<sup>b</sup> |
| **Refinement** |  |
| Amino-acid residues (chain A/B) | 253/252 |
| L-glutamate/BPAM344 | 2/2 |
| Chloride/sodium ions/water | 4/3/550 |
| *R*<sub>work</sub>/*R*<sub>free</sub> (%) | 15.3/17.9<sup>c</sup> |
| *Average B values (Å<sup>2</sup>) for:* |  |
| Amino-acid residues (chain A/B) | 19.2/21.3 |
| L-glutamate/BPAM344 | 9.5/11.1 |
| Chloride/sodium/water | 27.6/21.1/27.3 |
| RMSD bond length (Å)/angles (°) | 0.009/1.03 |
| Ramachandran outliers/favored (%) | 0/99.0<sup>d</sup> |
| Rotamer outliers (%) /Cβ outliers (%) /Clash score | 1.1/0/1.76<sup>d</sup> |

<sup>a</sup> a.u: asymmetric unit of the crystal
<sup>b</sup> Outershell values are shown in parentheses
<sup>c</sup> R<sub>free</sub> is equivalent to R<sub>work</sub>, but calculated with 5 % of reflections omitted from the refinement process
<sup>d</sup> MolProbity statistics from phenix

Distinct electron density (not shown) revealed the position of L-glutamate at the orthosteric site, positioned within the cleft between D1 and D2 (Fig. 1A). The binding interactions between L-glutamate and GluK2-LBD are similar to those previously reported for the structure of GluK2-LBD with L-glutamate only. Two BPAM344 molecules could be fitted at the allosteric sites at the dimer interface, based upon distinct electron density (Fig. 1C). One chloride ion and two sodium ions bind within the dimer interface at the top of the dimer (Fig. 1A), as previously reported [22, 23]. Chloride and sodium ions are necessary for the activation of KARs, thereby functioning as endogenous PAMs [24].

### Minor effect on domain closure

The D1-D2 domain closure of the structure of GluK2-LBD with L-glutamate and BPAM344 is 24.2° in chain A and 23.6° in chain B, with the domain closure of the GluK2 apo structure with BPAM344 set to zero (PDB ID: 8FQW; no ligand bound at the orthosteric site) [8] (Fig. 1B). For comparison, the domain closure of GluK2-LBD with L-glutamate only [10] is 23.1° in chain A and 22.3° in chain B (Fig. 2B). Thus, only a minor increase in the domain closure is observed upon addition of BPAM344. This suggests that the positive allosteric function of BPAM344 is not to increase domain closure in GluK2 but rather to stabilize the dimer interface, thereby delaying the entry of the receptor into the desensitized state, where the receptor is closed despite L-glutamate being present. This is in accordance with previous studies on LBD structures with PAMs [6, 25].

**Fig. 2.**
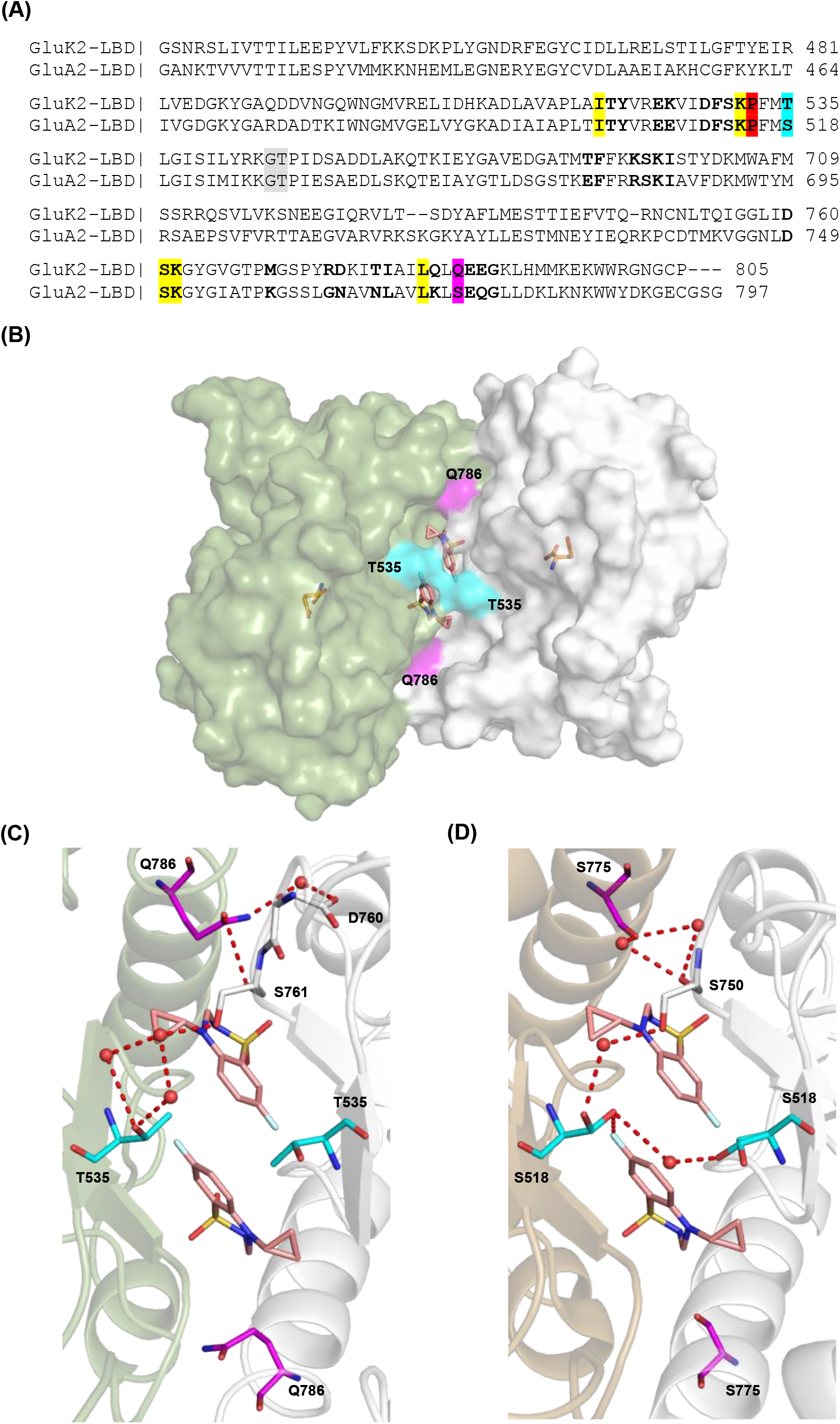
Comparison of structures of the kainate receptor GluK2-LBD and the AMPA receptor GluA2-LBD in the presence of L-glutamate and BPAM344. (A) Sequences alignment of LBDs from GluK2 and GluA2. Residues part of the dimer interface in GluK2-LBD are shown in bold as well as the corresponding residues in GluA2. Pro532 forming a direct hydrogen bond to BPAM344 in both receptor subtypes is shown in red. Dimer interface residues within 4 Å of BPAM344 that differ between GluK2 and GluA2 receptors are highlighted in cyan and magenta, respectively. Other residues within 4 Å of BPAM344 are highlighted in yellow. (B) GluK2-LBD dimer shown in surface representation with chain A in grey and chain A_sym_ in green. Thr535 and Gln786 are shown in surface representation and colored as in (A). The two BPAM344 molecules at the dimer interface and two L-glutamate molecules in the orthosteric sites located at the cleft between lobes D1 and D2 are shown in sticks representation with carbon atoms in salmon and orange, respectively. (C) Close-up on the binding site of BPAM344 in the GluK2-LBD complex with L-glutamate and BPAM344, showing direct and water-mediated contacts (red dashed lines) from Thr535 and Gln786, respectively, to the other subunit. (D) Close-up on the binding site of BPAM344 in GluA2-LBD with chain B in grey and chain C in sand, in complex with L-glutamate and BPAM344 (PDB ID: 4N07), showing direct and water-mediated contacts from Ser518 and Ser775, respectively, to the other subunit or BPAM344.

### Active-like conformation of the GluK2-LBD co-crystalized with L-glutamate and BPAM344

To investigate how BPAM344 affects the dimer interface, we measured distances at *i*) the top of the LBD (CA atom in Arg775 of chain A to CA atom in Arg775 of chain Asym), *ii*) the central D1-D2 hinge region where the two allosteric binding sites are allocated (Pro532), and *iii*) at the bottom of the LBD where the chain in the full-length receptor would continue into the linker region connecting to the TMD (Pro667). The Arg775-Arg775 and Pro532-Pro532 distances were measured to be 13.2 Å and 8.9 Å, respectively (Fig. 1B, top), which are similar to the GluK2-LBD structure with glutamate only (Fig. 1B, middle) and the full-length GluK2 apo structure with BPAM344 (Fig. 1B, bottom). The narrow space at the allosteric binding sites will limit the size of modulators being able to bind.

The Pro667-Pro667 distance in the structure of GluK2-LBD with L-glutamate and BPAM344 is 39.8 Å (Fig. 1B, top). In the AMPAR GluA2, the corresponding distance has previously been shown to correlate with the efficacy of agonists at AMPARs [26]. In the GluK2-LBD structure with L-glutamate only a 3 Å smaller distance of 36.7 Å is observed (Fig. 1B, middle), indicating a minor difference in the movement of lobe D2 upon binding of BPAM344. However, the distance is significantly decreased in the full-length GluK2 apo structure with BPAM344 (21.8 Å), in agreement with the receptor being in the resting state.

### Binding mode of BPAM344

A direct hydrogen bond (2.9 Å, chain A) is observed between the sulfonamide nitrogen atom in BPAM344 and the backbone oxygen atom of Pro532 (Fig. 1C). This direct hydrogen bond was also formed in the full-length structure of GluK2 apo with BPAM344 determined by cryo-EM [8] and in the GluK1-LBD crystal structure with kainate and BPAM344 [6]. In addition to the direct hydrogen bond, the binding of BPAM344 is mediated through numerous van der Waals interactions. Specifically, Ile519, Lys531, Pro532, Phe533, Met534, Thr535, Ser761, Lys762, Gly763, Leu783, Gln786, and Leu791 are found within 4 Å of BPAM344 (Fig. 1C). All these residues are conserved between GluK1 and GluK2. Together, the polar and nonpolar interactions help stabilizing the dimer interface of GluK2-LBD and thereby the active conformation of the receptor. The distance between the two BPAM344 molecules seems to be slightly smaller (4.1 Å) when L-glutamate is present compared to the apo structure (5.3 Å) (Fig. 1E). In comparison to the GluK2-LBD structure with L-glutamate only [10], the modulator displaces approximately eight water molecules within the dimer interface (not shown), which is similar to what was previously observed in GluK1-LBD [6]. In summary, the binding mode of BPAM344 in GluK2 is similar in the active-like and resting state conformations. Furthermore, the binding mode of BPAM344 in the active-like conformation is similar between GluK1 and GluK2.

### Molecular determinants for allosteric modulation by BPAM344

To address the molecular basis for the difference in potency of BPAM344 at GluK2 and GluA2, we focused on potential structural differences in the BPAM344 binding site and the surrounding dimer interface region between our structure and a previously determined structure of the GluA2-LBD in complex with BPAM344 and L-glutamate (PDB 4N07) [7]. Thirty-one interface residues in dimer A/Asym were identified using the PISA server [27], of which twelve differ between GluK2 and GluA2 (Fig. 2A). Considering residues within 4 Å of BPAM344, we identified eight residues among which only Thr535 in GluK2 (Ser518 in GluA2) and Gln786 in GluK2 (Ser775 in GluA2) differ (Fig. 2A-B).

In the GluA2-LBD, the hydroxyl group in Ser518 mediates a weak hydrogen bond to the fluorine atom in BPAM344 (Fig. 2D). In contrast, the hydroxyl group in Thr535 in GluK2 is not engaged in polar interaction with the modulator but points away from the fluorine atom in BPAM344 and forms water-mediated hydrogen bonds to Ser761 in the other subunit (Fig. 2C). In GluA2, the hydroxyl group in Ser518 may also create a polar contact to Ser518 of the other subunit through one water molecule (Fig. 2D). The side chain of Ser518 in GluA2 exists in two conformations with an occupancy of 0.33 and 0.46, respectively, for the two chains [7]. In the other conformation, one water molecule mediates a polar contact between Ser518 and Ser750 in the other subunit (Fig. 2D). In contrast, two water molecules are needed to generate the corresponding polar contact in GluK2 (between Thr535 and Ser761) as shown in Fig. 2C. It has previously been suggested that the ∼90-fold lower potency of BPAM344 at KARs compared to AMPARs may originate from unfavorable steric interaction with this threonine in KARs [6].

The second key residue differing between the two receptor families, Gln786 in GluK2, can either be Ser or Asn in GluA2 due to the formation of two isoforms: GluA2 flip (i) containing Ser775 and GluA2 flop (o) containing Asn775. BPAM344 has until date been crystallized in a flip-like isoform only [7]. These residues were not observed to form any direct or water-mediated hydrogen bonds to BPAM344. Instead, Gln786 forms a direct hydrogen bond to Ser761 and a water-mediated contact to Asp760 (Fig. 2C). The corresponding residue (Ser775) in GluA2 creates two water-mediated contacts to Ser750 in the other subunit (Fig. 2D). Taken together, both residues (Gln786 in GluK2 and Ser775 in GluA2) participate in the stabilization of the dimer interface and may therefore, indirectly contribute to the differences in potency of BPAM344.

### Assay for functional characterization of GluK2 T535S and Q786S mutants

To investigate the role of T535/Gln786 in GluK2 for potency of BPAM344, we substituted these residues with their GluA2 equivalents (Materials & Methods), individually and together to create the single point-mutants GluK2-T535S and GluK2-Q786S and the double-mutant GluK2-T535S-Q786S and expressed these by transient transfection in HEK293 cells for use in a fluorescence-based Ca^2+^ influx assay that we recently have developed for functional characterization of KAR and AMPAR receptor ligands in a 96-well plate-based format.

However, for characterization of PAMs, the previous use of the Ca^2+^ influx assay utilized stably receptor expressing cell lines [28–32]. Therefore, to validate the use of transiently transfected HEK293 cells in the assay, we determined the potency of BPAM344 at transiently expressed GluA2 in parallel with stably expressed GluA2(*Q*)i, and observed similar EC50 values (transient expression: 1.28 ± 0.36 µM, stable expression: 0.86 ± 0.19 µM; P = 0.292) that furthermore is similar to the previously determined EC50 value (0.90 ± 0.10 µM) determined on rat primary brain cultures [7] (Fig.3A).

**Fig. 3.**
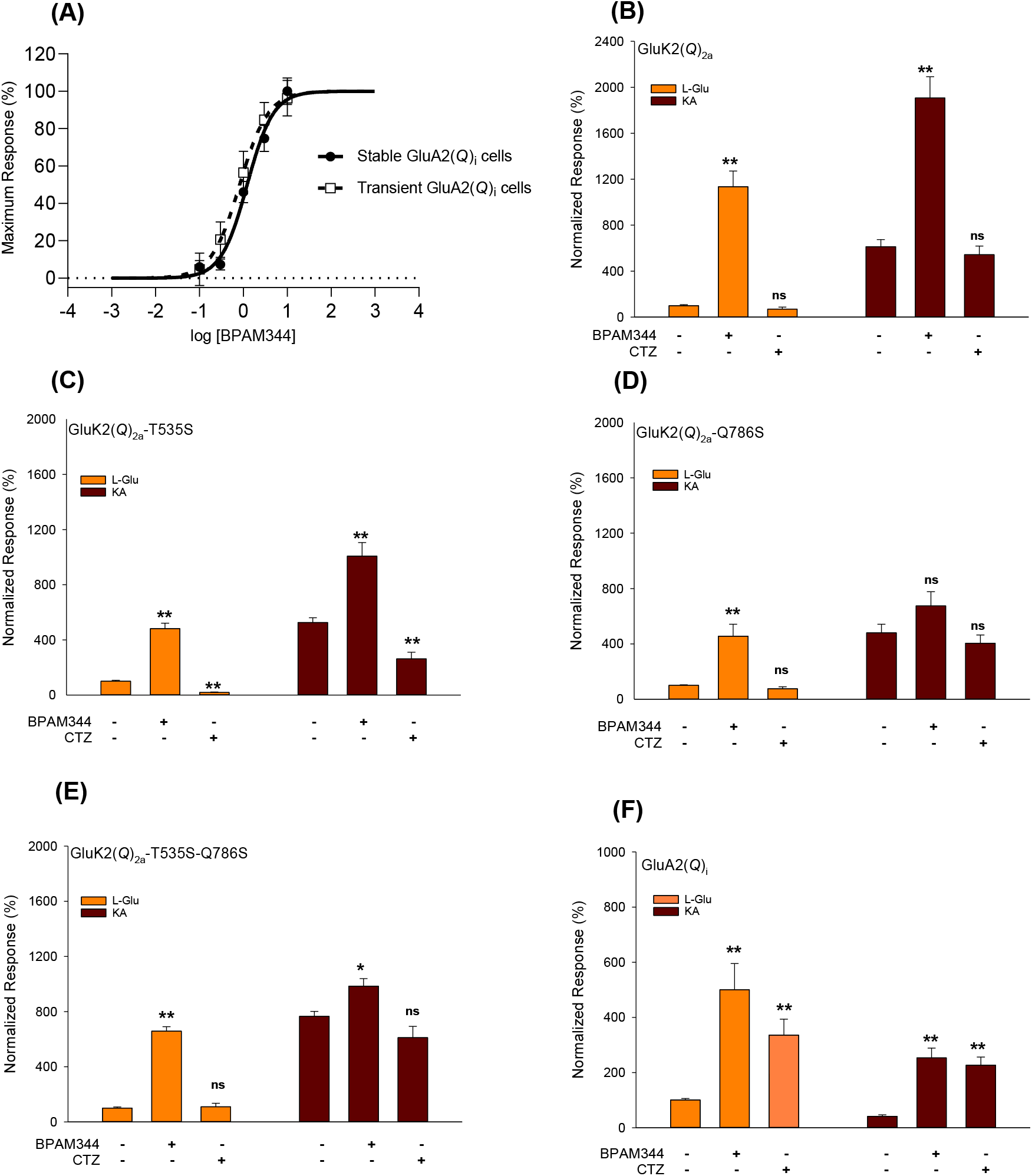
Functional characterization of WT GluA2, WT GluK2, and GluK2 mutants using a calcium-sensitive fluorescence-based assay, illustrating agonist and modulator preferences. (A) Assay validation test comparing the potency of BPAM344 at a stable GT-HEK293 GluA2(*Q*)_i_ cell line (unfilled circles) with transiently transfected GluA2(*Q*)_i_ GT-HEK293 cells (filled circles). Statistical analysis conducted in SigmaPlot with a Student t-test, showing no differences in the EC_50_ values (P value = 0.292). (B-F) Agonist and modulator preferences determined on WT GluK2(*Q*)_2a_ (B), mutant GluK2-T535S (C), GluK2-Q786S (D), GluK2-T535S-Q786S (E), and WT GluA2 (F), using a single concentration of 100 µM L-glutamate (L-Glu) or kainate (KA) with or without co-application of either BPAM344 or cyclothiazide (CTZ). The intracellular calcium response is normalized to the full response evoked from L-glutamate for all constructs. Bars are shown as means ± SEM of eight replicate wells presented as pooled data from three to four independent experiments.

**Fig. 4.**
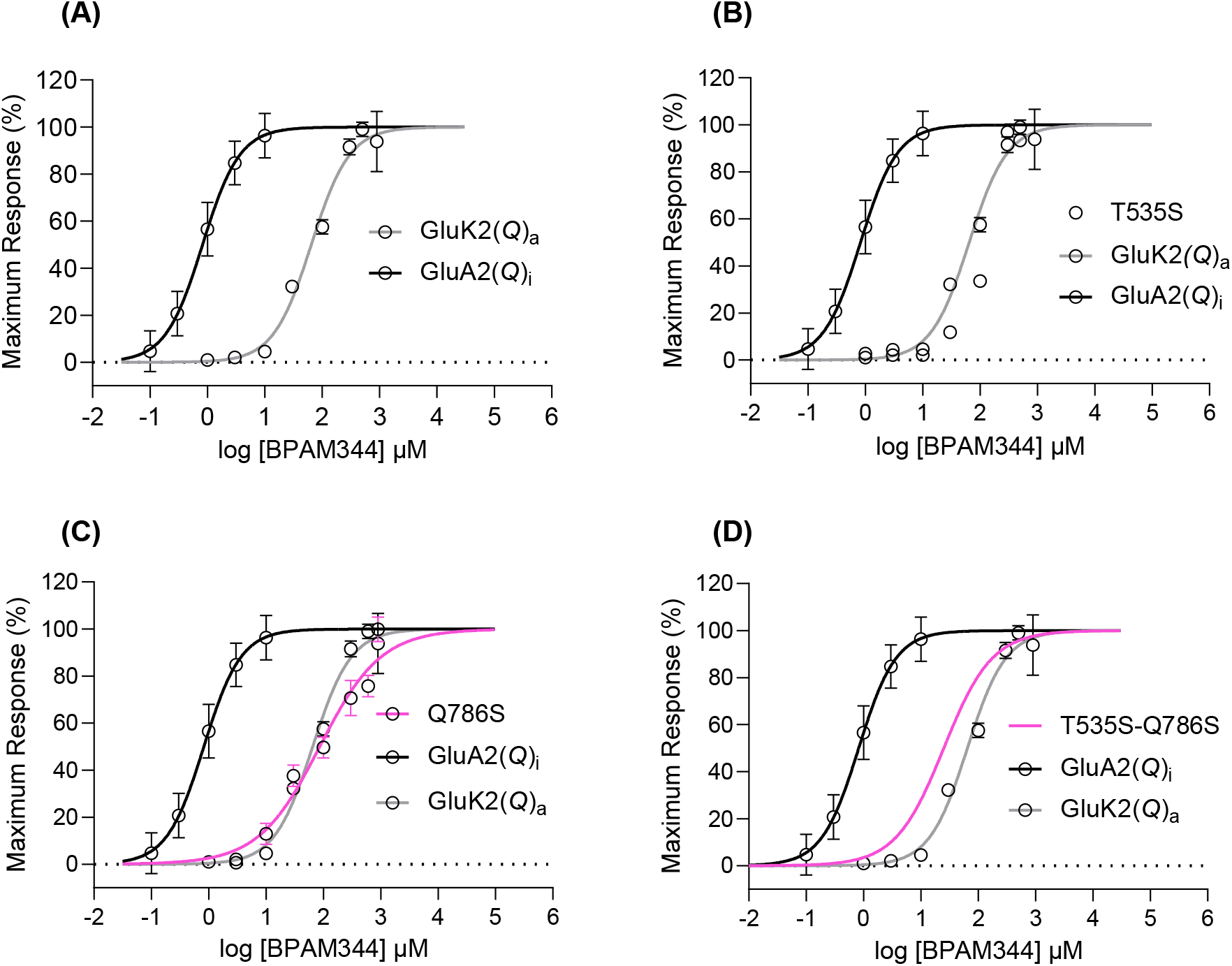
Dose-response curves from pooled experiments on BPAM344 at WT GluK2(*Q*)_2a_ and mutant GluK2-T535S, GluK2-Q786S, and GluK2-T535S-Q786S using the calcium-sensitive fluorescence-based assay and transiently expressed receptors. WT GluK2 (A), GluK2-T535S (B), GluK2-Q786S (C), and GluK2-T535S-Q786S (D). L-glutamate was used as agonist at 100 µM. EC_50_ values can be seen in Table 2.

### Agonist preference of BPAM344 at wild type and mutant receptors

Initially, we tested the response to L-glutamate and kainate of the T535S, Q786S, and T535S-Q786S GluK2 mutants in absence of potentiator and in presence of 100 µM BPAM344 or the AMPAR-selective potentiator cyclothiazide (CTZ) and compared the response profile to WT GluK2 and WT GluA2. In absence of potentiators, the WT GluK2 shows a higher response to kainate than L-glutamate (Fig. 3), whereas WT GluA2 shows smaller sized response to kainate than L-glutamate possibly due to kainate being a partial agonist at AMPARs [33].

Kainate acts as a more efficacious agonist at KARs which agrees with previous observations from calcium-sensitive fluorescence-based assays on KARs [28, 29]. All the mutants showed a response phenotype similar to WT GluK2 (Fig. 3, Table 2). Thus, the mutations in the BPAM344 binding site do not change the agonist profile of WT GluK2.

**Table 2.** Efficacy and potency of BPAM344 at WT GluA2(*Q*)_i_, WT GluK2(*Q*)_2a_, and mutants GluK2-T535S, GluK2-Q786, and GluK2-T535-Q786S. Utilizing the calcium-sensitive fluorescence-based assays and transiently expressed GT-HEK293 cells.

| Compounds | WT GluK2 | GluK2-T535S | GluK2-Q786S | GluK2-T535S-Q786S | WT GluA2 |
| --- | --- | --- | --- | --- | --- |
|  | % Response<sup>a</sup> | % Response<sup>a</sup> | % Response<sup>a</sup> | % Response<sup>a</sup> | % Response<sup>a</sup> |
| **L-Glutamate (L-Glu)** | 100 ± 9 | 100 ± 7 | 100 ± 7 | 100 ± 8 | 100 ± 6 |
| **L-Glu + BPAM344** | 1135 ± 137 | 482 ± 40 | 641 ± 193 | 658 ± 32 | 501 ± 95 |
| **L-Glu + CTZ** | 70 ± 18 | 19 ± 3 | 64 ± 18 | 109 ± 26 | 336 ± 58 |
| **Kainate (KA)** | 612 ± 62 | 526 ± 34 | 667 ± 114 | 765 ± 36 | 41 ± 5 |
| **KA + BPAM344** | 1907 ± 185 | 1008 ± 97 | 968 ± 204 | 984 ± 55 | 253 ± 35 |
| **KA + CTZ** | 543 ± 75 | 262 ± 49 | 578 ± 109 | 611 ± 82 | 227 ± 29 |
|  | Potency(μM) | Potency(μM) | Potency(μM) | Potency(μM) | Potency (μM) |
| **BPAM344** | (71.3 ± 5.2<sup>b</sup>) | (112 ± 13<sup>b</sup>) | (91.0 ± 10.4<sup>b</sup>) | (21.3 ± 3.9<sup>b</sup>) | (1.28 ± 0.36<sup>b</sup>) |

<sup>a</sup> Mean response relative to L-glutamate ± SEM (%). The data are obtained from 3-4 independent experiments performed in triplicate or quadruplicate wells for each compound.
<sup>b</sup> EC<sub>50</sub> value ± SEM was determined in the presence of 100 μM L-glutamate for GluA2(Q)<sub>i</sub>, GluK2(Q)<sub>2a</sub>, GluK2-T535S, GluK2-Q786S and GluK2-T535S-Q786S.

For WT GluK2, the response after application of BPAM344 was higher for L-glutamate (∼11-fold) than for kainate (∼3-fold). The same pattern was observed for the single mutants and the double mutant causing a 5-6-fold-potentiation in presence of L-glutamate but a 1.5-2-fold-potentiation with kainate (Fig. 3, Table 2). It would be interesting to attempt crystallization of GluK2-LBD with kainate, which, however, has not been successful in our hands. In contrast, the addition of BPAM344 led to a similar fold-potentiation (∼5-fold) of WT GluA2 regardless of agonist.

In summary, the choice of agonist seems to influence the response of BPAM344 at GluK2, both at WT GluK2 and in the presence of either threonine or serine at position 535 or glutamine or serine at position 786, whereas no preference is observed at WT GluA2.

### Cyclothiazide potentiates GluA2 but not GluK2 and mutants

Cyclothiazide (CTZ) is a well-studied positive allosteric modulator at AMPARs, evoking its action by binding to the dimer interface stabilizing the receptor [33]. CTZ binds in the same region in GluA2 as BPAM344 but adopts a different binding mode [25]. In contrast to BPAM344, CTZ has been reported to have no or minor antagonist effect on KAR currents evoked by L-glutamate or kainate [34, 35] or AMPARs containing a S775Q mutation [36, 37]. Therefore, we also tested CTZ at WT GluA2, WT GluK2, and at the three GluK2 mutants in our assay (Fig. 3, Table 2). Our results show that CTZ potentiates the currents of L-glutamate ∼3-fold at GluA2 and of kainate ∼2-fold, in accordance with the previous study concluding that CTZ acts as a PAM on AMPARs regardless of L-glutamate or kainate [38]. At WT GluK2, GluK2-T535S, GluK2-Q786S, and GluK2-T535S-Q786S, the co-application of CTZ with L-glutamate or kainate in all cases seem to cause a slight inhibition or no effect (Fig. 3B-E), also resembling previous observations [34, 35].

### BPAM344 potency increased at GluK2-T535S-Q786S

We then performed concentration-response experiments for the potentiating effect of BPAM344 to determine the potency (EC50) to see if introduction of the GluA2 residues into the binding pocket of GluK2 could increase the potency towards the EC50 observed at WT GluA2. Specifically, the potency of BPAM344 at WT GluK2 was ∼60-fold lower than at WT GluA2 (Table 2), similar to what has previously been observed [6, 7].

The potency of BPAM344 at GluK2-T535S was determined to 112 ± 13 µM and relatively higher than 71.3 ± 5.2 µM at WT GluK2, suggesting that the change from a Thr residue to the counter residue in WT GluA2 is unable to restore the higher potency of BPAM344 at GluK2. Of note, the counter residue in WT GluA2, Ser appears in two conformations in the structure, where only one conformation of the side chain can create the direct hydrogen bond to the fluorine atom of BPAM344. This indicates a weak hydrogen bond between Ser518 and BPAM344 in WT GluA2 which might explain why the GluK2-T535S mutation have minor impact on the potency of BPAM344.

When changing the Gln residue to Ser at position 786 in GluK2 receptors, the EC50 value increased to 91.0 ± 10.4 µM compared to 71.3 ± 5.2 µM at WT GluK2 (^ns^P value = 0.572). Thus, mutation of Gln into Ser at position 786 in GluK2 does not improve the potency of BPAM344 when compared to wild type. This seems in good agreement with the fact that BPAM344 forms no direct or water-mediated contacts to Gln786 in GluK2-LBD or Ser775 in GluA2-LBD.

In contrast to the single mutations, the double mutation GluK2-T535S-Q786S led to ∼3-fold improved potency from 71.0 ± 5.2 µM at WT GluK2 to 21.3 ± 3.9 µM at the double mutant (**P value =< 0.001). The substitutions may contribute to conformational changes in a way that either changes the kinetics of the ion channel or support the stability of the dimeric interface prolonging the duration of open ion channels by inhibition of desensitization.

However, introducing both mutations at the same time could not fully explain ∼60-fold less potency at GluK2 vs. GluA2 (Table 2). It is evident from previous studies [1] that the recovery from desensitization between AMPARs and KARs to L-glutamate is different with AMPARs displaying faster recovery than KARs, suggesting that the difference in potency of BPAM344 may also originate from the distinct desensitization present and could explain why a full recovery of BPAM344 potency was not observed when making WT GluK2 into a GluA2 alike receptor.

This new functional and structural insight into GluK2 and the binding mode BPAM344 may contribute to unravel the potential of KARs as targets for the treatment of brain diseases as well as aid in the search for tool compounds.

## Abbreviations

AMPA: α-amino-3-hydroxy-5-methylisoxazole-4-propionic acid
AMPAR: AMPA receptor
ATD: amino-terminal domain
BPAM344: 4-cyclopropyl-7-fluoro-3,4-dihydro-2*H*-benzo[*e*][1,2,4]thiadiazine 1,1-dioxide
cryo-EM: cryo-electron microscopy
CTD: carboxy-terminal domain
CTZ: cyclothiazide
EC50: half-maximal effective concentration
L-Glu: L-glutamate
GluK2-LBD: ligand-binding domain of GluK2
iGluRs: ionotropic glutamate receptors
KA: kainite
KARs: kainate receptors
LBD: ligand-binding domain
PAM: positive allosteric modulator
RFU: relative fluorescence unit
TMD: transmembrane domain
WT: wild type

## Acknowledgements

MAX-lab, Lund, Sweden, is thanked for providing beamtime and technical assistance. Linda Grønborg Dorvil is thanked for her technical support in the process of creating mutant constructs. Christophe Mulle is thanked for providing the GluK2 construct. Christina Kasper is thanked for providing the GluK2-LBD (S1S2) construct and Heidi Peterson for technical assistance in expressing and purifying GluK2-LBD protein. The Independent Research Fund Denmark – Medical Sciences (YB, JSK) and Danscatt (YB, MEJ, KF, JSK) are acknowledged for financial support. We acknowledge the MAX IV Laboratory for time on Beamline 911-3 under Proposal MX20130012. All funding sources had no part in the design of the study, the collection of data or in the analysis and interpretation, as well as no involvement in writing the manuscript or decision making of publication.

## Author Contributions

KF, DSP, ASK, and JSK conceived and supervised the study; YB, KF, ASK, and JSK designed experiments; YB and MEJ performed experiments; PF and BF synthesized the chemical compound BPAM344; YB, MEJ, KF, ASK, and JSK analyzed data; YB wrote the first draft of the manuscript; KF, ASK, and JSK made manuscript revisions. All authors commented and approved the final version of the manuscript.

## Data accessibility

The structure coordinates and corresponding structure factor files of the GluK2-LBD dimer with L-glutamate and BPAM344 have been deposited in the Protein Data Bank (PDB) under the accession code (PDB ID: XXXX).

